# Annotation Regression for Genome-Wide Association Studies with an Application to Psychiatric Genomic Consortium Data

**DOI:** 10.1101/049932

**Authors:** Sunyoung Shin, Sündüz Keleş

## Abstract

Although genome-wide association studies (GWAS) have been successful at finding thousands of disease-associated genetic variants (GVs), identifying causal variants and elucidating the mechanisms by which genotypes influence phenotypes are critical open questions. A key challenge is that a large percentage of disease-associated GVs are potential regulatory variants located in noncoding regions, making them difficult to interpret. Recent research efforts focus on going beyond annotating GVs by integrating functional annotation data with GWAS to prioritize GVs. However, applicability of these approaches is challenged by high dimensionality and heterogeneity of functional annotation data. Furthermore, existing methods often assume global associations of GVs with annotation data. This strong assumption is susceptible to violations for GVs involved in many complex diseases. To address these issues, we develop a general regression framework, named Annotation Regression for GWAS (ARoG). ARoG is based on finite mixture of linear regression models where GWAS association measures are viewed as responses and functional annotations as predictors. This mixture framework addresses heterogeneity of effects of GVs by grouping them into clusters and high dimensionality of the functional annotations by enabling annotation selection within each cluster. ARoG further employs permutation testing to evaluate the significance of selected annotations. Computational experiments indicate that ARoG can discover distinct associations between disease risk and functional annotations. Application of ARoG to autism and schizophrenia data from Psychiatric Genomics Consortium led to identification of GVs that significantly affect interactions of several transcription factors with DNA as potential mechanisms contributing to these disorders.

## 1 Introduction

Although genome-wide association studies (GWAS) have successfully identified thousands of genetic loci associated with human diseases, the design and analysis of these studies are challenged in two critical aspects. First, existing GWAS have revealed that, for many common disorders, the typical genetic architecture encompasses many genetic variants (GV) with individually small effects on phenotypes [29], indicating the need for larger sample sizes to reliably identify them. Second, roles of a large proportion of identified GVs remain elusive since they reside in non-coding sequences within introns or in regions between genes. Today, with the availability of affordable whole genome sequencing, our ability to elucidate the mechanisms by which genotypes influence phenotypes is far behind our ability to identify phenotype-associated variants.

In parallel to the rapid developments in the design and analysis of GWAS, large consortia projects such as the Encyclopedia of DNA Elements (ENCODE [13, 35]), the Roadmap Epigenomics Mapping Consortium (REMC [23]), the Genotype-Tissue Expression Project (GTEx [30]), and the International Human Epigenome Consortium (IHEC [1]) as well as many investigator-driven projects are generating diverse data types of RNA transcription (RNA-seq), DNA accessibility (DNase-seq), DNA methylation (Methyl-seq), protein-DNA interactions (ChIP-seq/exo), protein-RNA interactions (CLIP-seq), and chromatin state (Histone ChIP-seq) across diverse cell/tissue types. There is a growing literature on methods for utilizing one or more classes of these functional annotation data to support GWAS results. In contrast to methods requiring individual level GWAS data which are neither available publicly nor immediately [14, 33], many existing methods have a useful key feature of using population level data in the form of summary statistics of GVs [9, 12, 16, 31]. However, annotation data have rarely played more than an indirect role in assessing evidence for association in both types of approaches. Specifically, Iversen et al. [14], Pickrell [20] and Thompson et al. [31] used annotations to model prior probabilities of association of GVs under Bayesian framework or hierarchical modeling. Chung et al. [9] and Kichaev et al. [16] developed models to integrate binarized functional annotation data and GWAS summary statistics such as p-values and z-scores of GVs. Gagliano et al. [12] correlated annotations with GWAS association status, and used Bayes factor for the annotations to estimate the posterior odds of association of GVs for prioritization. A significant shortcoming of these methods is that they aim to globally relate associations of GVs to functional annotation data despite the fact that the same disease mechanism might be governed by distinct functional annotations. For example, disruption of an important pathway may arise by GVs in coding regions of the genes and/or in their regulatory mechanisms. Regulatory GVs may have a variety of mechanisms such as transcription factor (TF) binding, histone modifications, enhancer activity through chromatin architecture, DNA methylation, and alternative splicing [19]. Another key shortcoming is that they pre-select relevant annotations or use conveniently available annotations ignoring disease etiology, because they are not equipped to automatically select important annotations in a data-adaptive manner. Furthermore, several of them can only use annotation data in specific formats (e.g., most recent genetic analysis incorporating pleiotropy and annotation (GPA) [9] requires binary annotation variables).

To overcome these challenges, we develop a regression framework named Annotation Regression for GWAS (ARoG) and integrate GWAS and functional annotation data. ARoG models GWAS association measures, e.g., z-scores from univariate analysis of GWAS, as a linear function of functional annotations. It employs a mixture of linear regressions framework to accommodate the heterogeneity of associations between GWAS association measures and functional annotations. It aims to capture locally distinct associations that would not be revealed with an analysis that assumes homogeneity of these associations. A critical aspect of ARoG is that it can automatically select relevant annotations among a large number of annotations with penalization techniques. ARoG accommodates both categorical or continuous annotation types, both of which are commonly available. The rest of the paper is organized as follows. Section 2 presents empirical observations regarding GWAS association measures and functional annotations using Psychiatric Genomics Consortium (PGC) data. Section 3 develops ARoG and discusses implementation details. In Section 4, we analyze PGC autism and schizophrenia data and identify GVs that might influence these diseases with the potential to modulate TF-DNA interactions. Section 5 presents computational experiments with a wide variety of settings including PGC analysis-driven simulations. In Section 6, we provide concluding remarks and discuss extensions.

## 2 Exploring Psychiatric Genomics Consortium Data with Functional Annotations

PGC has conducted mega analysis of GWAS data for five psychiatric disorders: attention deficit/hyperactivity disorder (ADHD), autism spectrum disorder (AUT), bipolar disorder (BIP), major depressive disorder (MDD), and schizophrenia (SCZ) [2, 10]. They identified 4 genome-wide significant loci for BIP [21], and more than 100 genome-wide significant loci for SCZ [24, 25]. However, their analysis did not lead to any reproducible genome-wide significant loci for ADHD and MDD, and the analysis on AUT is in progress. Their GWAS summary datasets are publicly available at http://www.med.unc.edu/pgc/downloads. In what follows, we focus on AUT and SCZ data.

### 2.1 Autism GWAS

We generated a set of candidate SNPs by starting with the intersection of SNPs genotyped in all five disorder datasets from the PGC cross-disorder study [10]. After lifting the original genomic coordinates from hg18 to hg19, we obtained 1,219,561 SNPs common to all five disorders. We next selected a subset of the SNPs with a Benjamini-Hochberg (BH) adjusted association p-value smaller than or equal to 0.1 in any of the five disorders [6]. This led to a total of 1,430 SNPs. Next, we included 761 linkage disequilibrium (LD) partners of these SNPs as identified by the SNAP tool [15] with an *r*^2^ ≥ 0.8 to one or more of the 1, 430 SNPs. As part of pre-processing, we discarded SNPs with more than one reference allele, SNPs with nucleotide mismatches between the PGC dataset and the SNP database dbSNP [4], and SNPs not listed in dbSNP [4]. As a result, we obtained a total of 2,191 SNPs for analysis. Supplementary Figure 1(a) displays the histogram of the autism z-scores for these sets of SNPs and illustrates that, as expected, LD partners tend to contribute z-scores around zero to the overall distribution since they had BH adjusted p-values larger than 0.1 in the initial selection step. The manhattan plot of the p-values in Figure 1(a) indicates that the SNPs with the strongest association are on chr 5 (6 of them) and chr 6 (3 of them). All of these have raw p-values less than 10^−6^; however, they make neither the conventional GWAS p-value cutoff of 5 × 10^−8^ nor the Bonferroni cutoff of 4.1 × 10^−8^ specific for this study, indicating that common practice for GWAS analysis would not confidently identify significant SNPs from this study.

Currently, most integrative analysis methods consider functional annotations enabled by the large scale analysis results of consortia projects such as ENCODE (e.g., [9, 16, 20]). In our exposition, we consider a class of functional annotation which computationally quantifies effects of SNPs on TF binding. We used atSNP [37] for this quantification and created an annotation score matrix for the 2,191 SNPs [37]. Specifically, atSNP computes the likelihood that a given SNP disrupts or enhances the binding sites from a given set of position weight matrices (PWMs) characterizing the class of sequences which TFs recognize [28]. atSNP scans through subsequences overlapping with the SNP position with reference and SNP alleles for the best matches of both to a given PWM and quantifies the significance of the best matches with both alleles by p-values. The natural logarithm of the ratio of the two p-values is defined as the atSNP annotation score, which empirically reflects the change in the ranks of the PWM matches of the alleles. SNPs likely to enhance or disrupt binding of given TF have large absolute atSNP scores for the corresponding PWM while SNPs with little potential impact on binding have scores close to zero. We refer to Zuo et al. [37] for further computational details. We considered the JASPAR CORE database [17] for vertebrates with 205 PWMs as our motif library and scored the SNP set. Supplementary Figure 1(b) displays the heatmap of the resulting annotation score matrix along with the z-scores. Here, most SNPs have relatively weak annotation scores and only SNPs colored as dark green or red are likely to lead to significant changes (as assessed by atSNP p-values) in TF binding. As part of our exploratory analysis, we first regressed z-scores from each of the five disorders on each annotation score separately. Figure 1(b) displays the-log10 transformed p-values from these marginal regressions. We note that the overall association of the z-scores and functional annotations for some diseases are apparent (e.g., MDD). However, for autism, none of the annotations can be deemed as contributing to the variation in the autism z-scores based on this global marginal analysis as all BH adjusted p-values are greater than 0.1 (Supplementary Figure 1(c)).

**Fig. 1:**
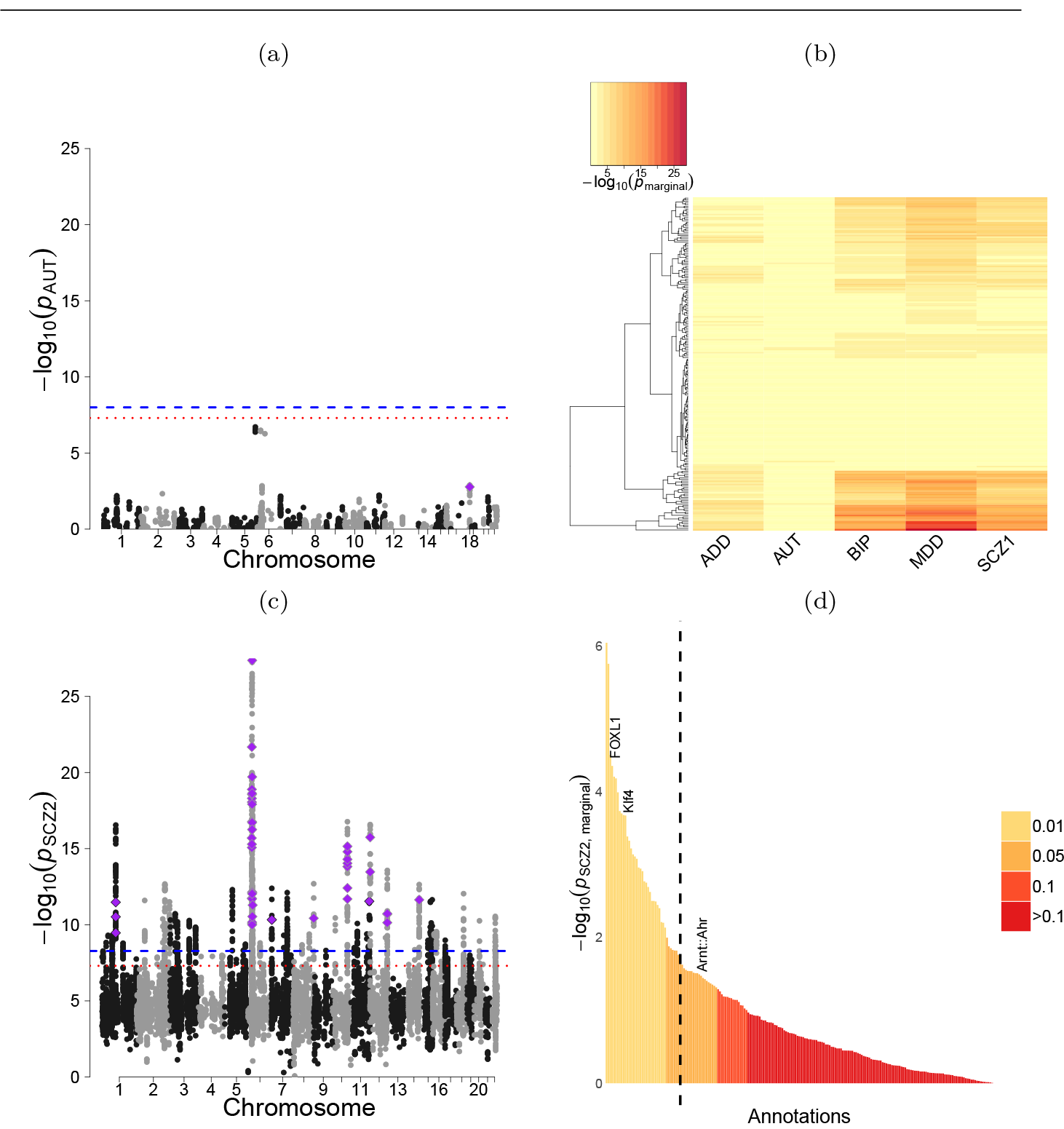
(a) Manhattan plot for autism association p-values across all 2,191 SNPs. *ARoG SNP* identified in Section 4.1 is marked with a purple diamond. Blue dotted and red dashed horizontal lines depict the Bonferroni cut-off at significance level of 0.05 and the conventional p-value cutoff of 5 × 10^−8^. (b) Heatmap of-log10 p-values from marginal regressions of z-scores for five psychiatric disorders on annotation scores. (c) Manhattan plot for SCZ2 association p-values across all 11,386 SNPs. *ARoG SNPs* identified in Section 4.3 are marked with purple diamonds. Blue dotted and red dashed horizontal lines depict the Bonferroni cut-off at significance level of 0.05 and the conventional p-value cutoff of 5 × 10^−8^. (d) Ranking of annotations based on marginal regressions of the SCZ2 z-scores on annotation scores for each TF. FOXL1, Klf4, and Arnt::Ahr TFs that are identified as associated with the z-scores in Section 4.3 are labeled. The dashed vertical line depicts the BH cut-off at significance level of 0.1.

### 2.2 Schizophrenia GWAS

PGC provides analyses of two schizophrenia GWAS: a SCZ study from [24] and a more comprehensive SCZ study from [25]. We refer to the first study as SCZ1 and the second study as SCZ2. SCZ1 and SCZ2 have genotypes for 1,252,901 and 9,444,230 SNPs, respectively. We considered 1,179,262 SNPs common to SCZ1 and SCZ2, filtered out SNPs with BH adjusted p-values larger than 0.01 for both studies, and retained the remaining 8,029 SNPs. Similar to the autism analysis, we excluded SNPs with multiple reference alleles or mismatches of alleles between the PGC datasets and dbSNP, and SNPs that are not in dbSNP. We next extended this set by including their LD partners with *r*^2^ ≥0.8. Our final set of SNPs for the analysis included 11,386 SNPs. z-scores of these SNPs have bimodal distributions in both SCZ1 and SCZ2 (Supplementary Figure 1(d)). Long tails of the z-score distribution of SCZ2 indicate many more statistically significant SNPs from this study. This may imply increased precision of SCZ2 over SCZ1, which is attributable to the seven-fold increase in the sample size. The manhattan plot in Figure 1(c) indicates that genome-wide significant SNPs from SCZ2 spread throughout the genome. In what follows, we used SCZ2 as the main analysis dataset for both marginal and ARoG analyses and SCZ1 as the validation dataset.

We generated an 11, 386 × 205 annotation score matrix using atSNP with the JASPAR PWM library. Supplementary Figure 1(e) displays the heatmap of the resulting annotation score matrix along with the SCZ1 and SCZ2 z-scores and illustrates that only a small proportion of SNPs might impact binding of a small subset of TFs. Marginal regressions of SCZ2 z-scores on annotation scores identify 40 annotations which are significant when adjusted for multiple testing by the BH procedure at level 0.1 (Figure 1(d)). However, given the large sample size, i.e., the number of SNPs, we view the associations from this marginal analysis as suggestive and turn our attention to developing a framework to identify subgroups of SNPs whose association measures can be explained by a subgroup of functional annotations.

## 3 A Mixture of Linear Regressions Framework for Incorporating Functional Annotations into GWAS Analysis

ARoG utilizes association measures of SNPs with disease/phenotype as response and potential effects of SNPs on TF binding as predictors as in Section 2. It exploits these functional annotations in a regression framework and aims to simultaneously boost detection power of SNPs and select relevant annotations.

### 3.1 Basic Annotation Regression for GWAS (ARoG(I))

Let *z_i_* ∈ℝ, =1, ………, n denote z-scores for n SNPs from a GWAS. Let X=[X*i*]*i*=1,………, *_n_* ∈ℝ^n×(*p*+1)^ denote an annotation score matrix for the SNPs, where X_*i*_=(X_*i*0_, ………, *X_ip_*) ∈ ℝ^*P*+1^ is a vector of *p* functional annotations for the i-th SNP with the first element of 1 as the intercept term. ARoG assumes that *n* SNPs are partitioned into *K* clusters and uses finite mixture of linear regression models (FMR) to relate the response *z_i_* and the predictor vector *X_i_* for SNP *i*, = 1,………, *n*. Basic ARoG, denoted by ARoG(I), performs *l*_1_-norm penalized maximum likelihood estimation for FMR (FMRLasso) proposed in Stadler et al. [27]. Following the notation of FMRLasso, we denote the prior probability of the *k*-th cluster as *π_k_*, its regression parameters as *β_k_* = (*β,_ko_*,………, *β_kP_*)^T^, and its variance as 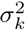. The conditional density function of z-scores, that is *z* given a functional annotation vector, *X* is then

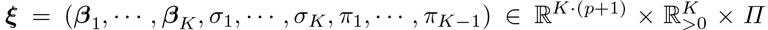

where 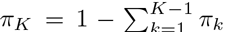 and 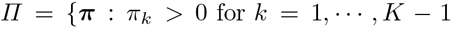 and 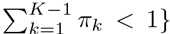 is a parameter space of π with 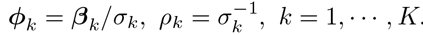. Here, β*_k_*;are cluster-specific regression parameters specifying how z-scores relate to the annotation scores within cluster *k*. Städler et al. [27] considered a reparametrized form of this density for scale-invariant estimation and efficient computation. Specifically, they reparametrized the regression parameters and the variances as follows:

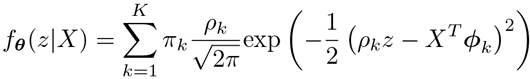

We can rewrite equation (1) with the new parameters as

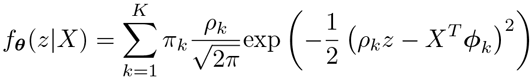

where 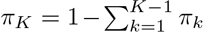 is a new parameter vector and Π is the same set as above with 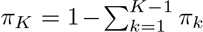

FMRLasso penalizes the negative log-likelihood with an *1*_1_ norm penalt [32]:

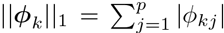

where μ is a tuning parameter and 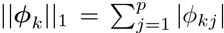. The penalty term weighs the contributions from each cluster by the corresponding prior probabilities. For a given number of clusters, *K*, and a given tuning parameter, μ, we define a minimizer of the penalized negative log likelihood as the FMRLasso estimator, denoted by 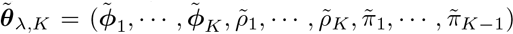 Städler et al. [27] also suggested an unweighted penalty term ∑_*k*_||ϕ_*k*_||_1_ and another weighted penalty term of the form 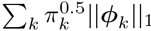. The unweighted penalty term tends to perform poorly in unbalanced cases, where the numbers of SNPs across clusters differ significantly [27]. Therefore, ARoG utilizes the weighted penalty term with *π_k_*, which performs well in both the balanced and unbalanced cases.

A key issue in the mixture linear regression model is the selection of the optimal number of clusters and the optimal tuning parameter. We use a modified Bayesian Information Criteria (BIC), defined by Städler et al. [27] as

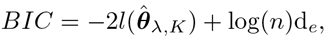

where 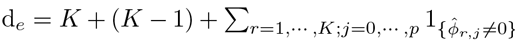 is the effective number of parameters. We perform a grid search over a set of (μ, K) and find the optimal combinations, 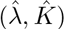, achieving the smallest modified BIC. Städler et al. [27] showed that a single cluster model selects no variables with 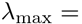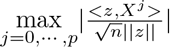, where *X^j^* is the (*j*+1)-th column of **X**, and suggested λ_max_ as the upper bound for the value of the tuning parameter. ARoG increases this upper bound three to six times since multiple clusters may require a larger tuning parameter to avoid selecting false positive annotations. With a slight abuse of notation, we denote the ARoG(I) parameter estimates as 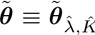. The annotation coefficient and the variance for the *k*th cluster are estimated by 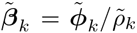 and 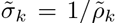, respectively. The posterior probability that SNP *i* belongs to the *k*th cluster is given by

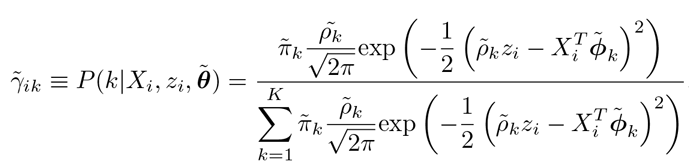

ARoG assigns the SNPs to the clusters for which they have the largest posterior probabilities. This generates K SNP sets with members 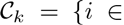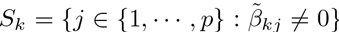 and set sizes |C_*k*_| = *n_k_*, *k* = 1, ………, *K*. We denote the selected annotation set of each cluster as 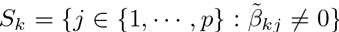 with numbers of the selected annotations, |S*_k_*| = *p_k_* ≤ *p*, and define the entire annotations selected by the model as the union of the selected annotations across all clusters, *S* = ∪*_k_S_k_* C { 1, ………, *p* }.

### 3.2 Permutation Testing for ARoG

The ARoG framework follows up the penalized likelihood-based selection with a permutation testing to evaluate the significance of the selected annotations, which typically have small effect sizes based on our data analysis results in Section 4. We specifically test whether the maximum absolute value of the coefficients of each functional annotation across all clusters can arise by chance.We randomly permute the z-scores of the SNPs a large number of times (at least 1000 times), and refit ARoG to each permuted dataset. At each fit,we record the maximum absolute value of the estimated coefficients for each annotation across all clusters, 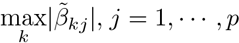. This collection generates functional annotation specific null distributions. Then the p-value for the *j*-th annotation is computed as the proportion of datasets with the maximum absolute value of the estimated coefficients for the annotation larger than 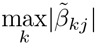. We utilize the BH procedure [6] at level 0.1 to account for the multiplicity of the annotations.

### 3.3 Two-step Annotation Regression for GWAS (ARoG(II))

Basic ARoG filters false positive annotations with a global penalization across all clusters; however, it is still prone to selecting a nonignorable number of false positive variables as both the simulations of Städler et al. [27] and our computational experiments in Section 5 illustrate. To reduce this effect and, thereby, increase specificity, we propose and study two-step ARoG, denoted by ARoG(II). ARoG(II) implements a cluster level penalization and a refit estimation after the initial global penalization by ARoG(I). The additional cluster level penalization is similar to relaxed Lasso of Meinshausen [18] which employs another level of Lasso in the context of standard multivariate linear regression model. Both ARoG(II) and relaxed Lasso aim to filter out false positive variables resulting from the initial penalization and thereby lead to better or comparable prediction with more accurate variable selection. Refitting has been widely used as a simple but practical tool to overcome the biased estimation of Lasso [8]. The refit step aims to improve regression parameter estimation by alleviating shrinkage effects towards zero due to penalization.

ARoG(II) starts with the optimal clusters selected by the initial FMRLasso of ARoG(I), and considers a standard multivariate linear regression model for each cluster with its FMRLasso selected annotation set. Specifically, ARoG(II) idds an *1*_1_ penalty term to the residual sum of squares within each cluster, hus obtains the following preliminary estimators:

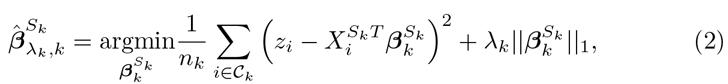

where 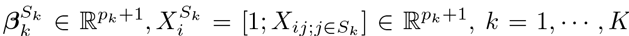. The tuning parameter selection for each cluster is through BIC [26] and the coefficients estimated with the optimal tuning parameter are denoted as 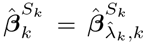.Next, based on the clusterwise Lasso, we obtain a smaller annotation set, 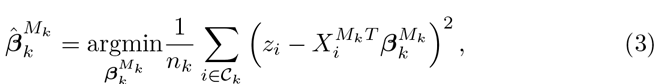 with size |*M_k_*| = *d_k_* ≤ *p_k_*, and refit a least squares regression with this annotation set:

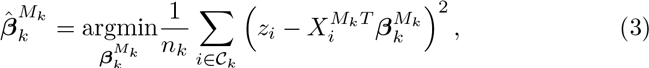

where 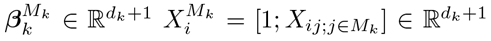. We then have ARoG(II) annotation score coefficients for the k-th cluster as

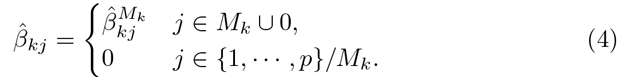

Similar to ARUG(I), we define AROG(II) annotations as the union of selected annotations over the clusters, *M* = ∪ *_k_M_k_* ⊆ *S*. There is a trade-off between ARoG(I) and ARoG(II) since cluster-level Lasso tends to gain specificity and lose sensitivity with more aggressive annotation screening. The level of the trade-off varies on a case by case basis. We further discuss this issue with computational experiments in Section 5.1. The permutation testing described in Section 3.2 is also part of ARoG(II).

### 3.4 Numerical Implementation

We implement ARoG with publicly available R packages `fmrlasso` and `glmnet`. The `fmrlasso` package fits FMRLasso with a block coordinate descent generalized expectation-maximization algorithm (BCD-GEM) proposed by Städler et al. [27]. It alternates between an expectation step (E-step) and a generalized maximization step (generalized M-step), which updates the prior probabilities, π at once, then updates the reparametrized regression coefficients, ϕ and standard deviations, ρ. In the M-step, coordinate updates for ϕ and ρ are performed on decoupled K optimization problems for each cluster separately. We use a hierarchical clustering strategy to initialize the BCD-GEM algorithm. This clustering operates on a distance matrix where the distance between any two SNPs is calculated as *l*_1_ distance of their z-scores and summary annotation scores, which is defined as the *l*_2_ norm on the annotation score vector. This distance criterion ensures that SNPs with similar z-scores and similar variability in the functional annotations are more close to each other. As a result of this clustering each SNP has initial membership probabilities for the cluster it belongs to and the remaining clusters with a ratio of 9 to i. For the M-step, we initialize 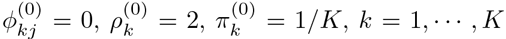 and *j* = ………,*p* Finally, we implement the cluster level Lasso of ARoG(II) with a coordinate descent algorithm using glmnet.

## 4 ARoG Analysis of PGC Data

### 4.1 PGC Autism GWAS

We fitted ARoG(I) and ARoG(II) to the autism dataset described in Section 2.i and varied the number of clusters as K = 1, ………, 10. Table i presents parameter estimates from both ARoGs with 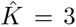 as the optimal number of clusters. Refitting for ARoG(II) is performed after each SNP is assigned to the cluster for which the SNP has the highest posterior probability based on ARoG(I) to reestimate both the regression parameters and the cluster-specific variances. Both ARoGs have the first and the second clusters as intercept-only models and select FOXLi and Nkx2-5 TFs for the third cluster. We kept the ARoG(I) intercept estimate for the first cluster since no SNPs were assigned to this cluster. Estimated coefficients for both TFs of the cluster 3 indicate that the SNP-driven increase in binding affinities for FOXLi and Nkx2-5 associate with the increase in autism risk in cluster 3. We further support the significance of these associations with a permutation test described in Section 3.2 (Supplementary Figure 2(a)). The third cluster has a total of thirteen SNPs, nine of which constitute the most genome-wide significant SNPs depicted in the Manhattan plot of Figure i(a). As an alternative multivariate approach to ARoG, we also used ordinary least squares (OLS) and Lasso regression to select the most relevant annotations from the set of 205. OLS did not select any annotations with a BH adjustment on the OLS p-values at level 0.i and had unadjusted p-values of 0.332 and 0.0i4 for FOXLi and Nkx2-5, respectively. We obtained 11 Lasso-selected annotations including Nkx2-5 and FOXLi with 5-fold cross-validation to tune the *l*_1_ penalty parameter. However, neither of these survived the permutation testing implemented in a way similar to that of ARoG’s (Supplementary Figure 2(a)). This analysis suggests that ARoG is indeed exploiting associations detectable only when appropriate subgroups of SNPs are considered.

We investigated the effects of the selected annotations, FOXLi and Nkx2-5, on autism z-scores of cluster 3. Figure 2(a) highlights significant positive associations of the z-scores with FOXLi and Nkx2-5 annotation scores, within cluster 3. Considering the whole set of SNPs leads to weak positive associations without statistical support from marginal regressions for both TFs. Figure 2(b) displays the heatmap of FOXLi and Nkx2-5 annotation scores of cluster 3 SNPs organized by hierarchical clustering along with their z-scores on top and supports that the variation in z-scores is well explained by these two annotation scores.

**Fig. 2:**
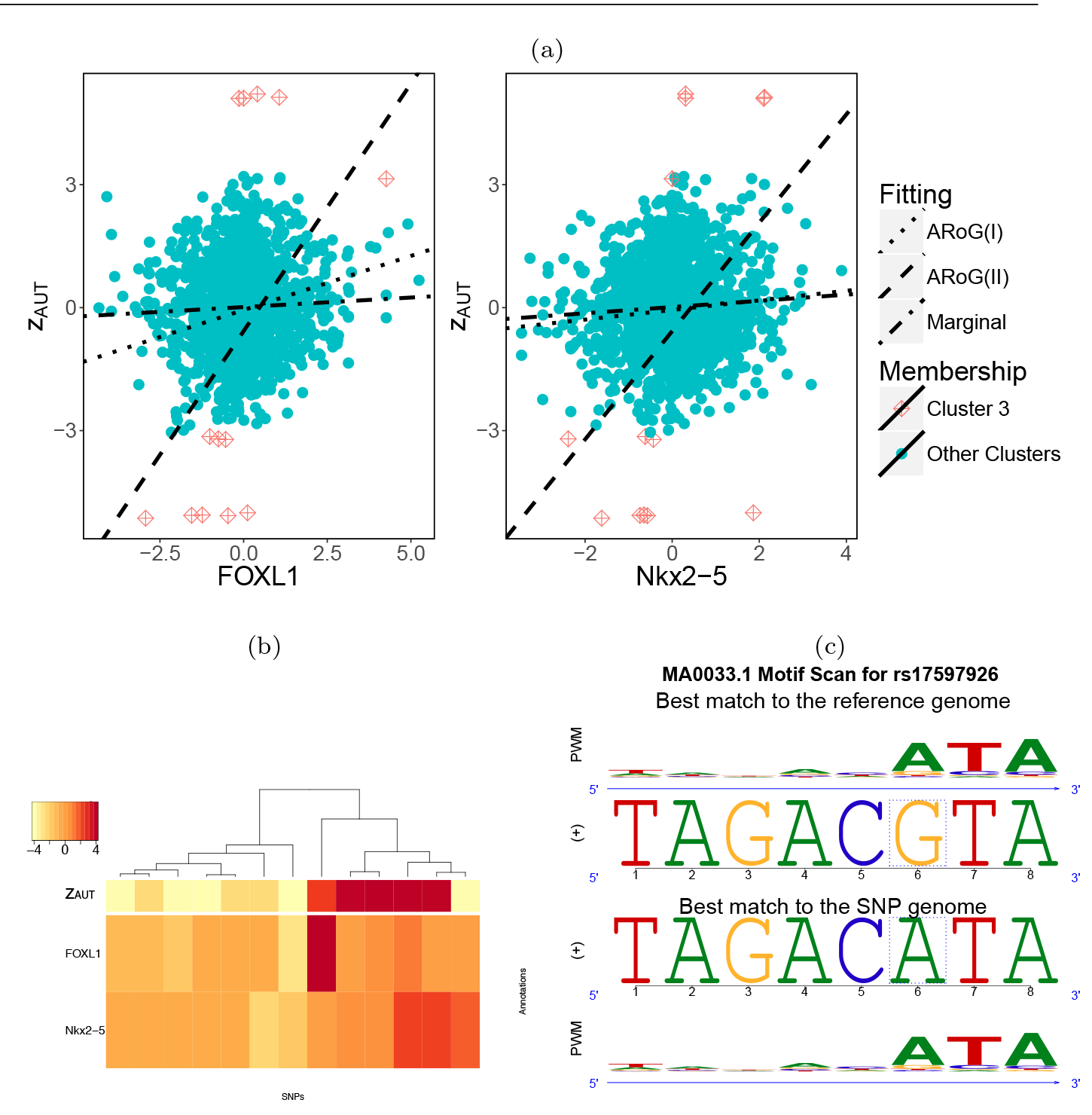
((a) Autism z-scores vs. annotation scores for FOXL1 and Nkx2-5 TFs selected for cluster 3 along with the marginal linear regression line fit and ARoG estimates (intercept and slopes for FOXL1 and Nkx2-5). (b) Hierarchical clustering of SNPs in Cluster 3 based on ARoG selected annotations with their AUT z-scores. (c) Composite sequence logo of SNP rs17597926 with the FOXL1 PWM. The middle two rows represent best matching genomic subsequences to the FOXL1 PWM with the reference and SNP rs17597926 alleles respectively. The dashed boxes mark the SNP location. Top and bottom rows display FOXL1 PWM sequence logos aligned to the best reference and SNP allele matches.

**Table 1:**
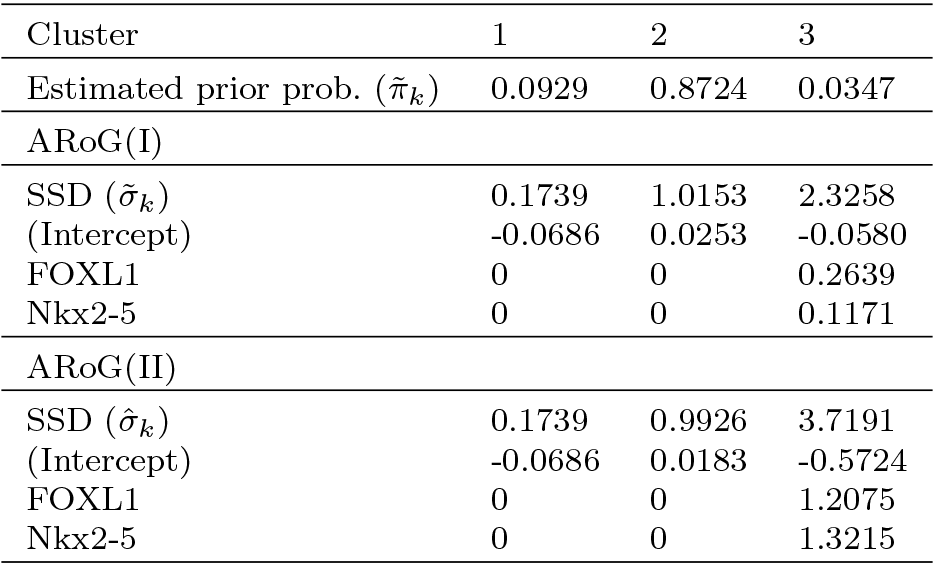
ARoG parameter estimates with PGC autism data.

Next, we define ARoG driven candidate causal SNPs, i.e., *ARoG SNPs*, as SNPs leading to significant TF binding affinity changes and having marginal association with the disorder. atSNP reports p-values assessing whether the observed change in TF binding affinity due to SNP is significant. We use these p-values along with raw GWAS association p-values to refine the SNPs in cluster 3 and create a set of *ARoG SNPs*. For the autism application, we considered a subset of the cluster 3 SNPs with raw GWAS p-value of at most 0.005 and atSNP p-value of at most 0.01 for FOXL1 or Nkx2-5, resulting in a single *ARoG SNP*, rs17597926. The unadjusted p-value of this SNP from autism GWAS is 0.0017 and the resulting atSNP FOXL1 p-value is 0.0008. The composite logo plot in Figure 2(c) confirms that rs17597926 is creating a potential FOXL1 binding site. rs17597926 is located within the 5th intron of the TCF4 gene, known to interact with helix-loop-helix proteins and regulate neurodevelopment [11]. Furthermore, this SNP has been identified as a *cis*- eQTL for TCF4 in a recent brain expression GWAS [36]. This is an additional support for a potential regulatory role of rs17597926 as a mediator of TCF4 gene in psychiatric disorders.

### 4.2 Comparison with GPA on PGC Autism GWAS

In addition to the ARoG analysis, we also applied the GPA approach of Chung et al. [9] to the autism data. We emphasize that GPA and ARoG approaches utilize functional annotations from different angles: GPA goes after global signals using all SNPs genotyped whereas ARoG aims to identify local signals by focusing on a smaller set of signals with potential significance. GPA is based on a joint generative model of association p-values of the SNPs and annotation data and identifies annotations that the disease-associated SNPs are enriched for. It aims to simultaneously identify null (SNPs not associated with the phenotype) and non-null (SNPs associated with the phenotype) and quantify the enrichment of a given annotation within these SNP sets. It specifically tests whether equal proportions of non-null and null SNPs carry the annotation. Although it can handle multiple annotations simultaneously, our results from two application schemes of “one annotation at a time” versus “all annotations simultaneously” showed extreme differences which could potentially be attributable to the violation of the GPA independence assumption of the annotations given the SNPs null versus non-null status. As a result, we focus on applying GPA one annotation at a time. Of the 1,219,561 genotyped SNPs in the PGC cross-disorder study, we considered a subset of 1,210,570 SNPs that were also in the dbSNP database. GPA works with binary annotations; therefore, we created a binary annotation score matrix by running atSNP [37] on the SNPs and thresholding atSNP p-values at 0.05. We applied GPA to each of the 205 annotations separately and estimated the proportions of null and non-null SNPs associated with each annotation. Results of GPA hypothesis testing for annotation enrichment did not identify any annotation as significantly enriched for autism-associated SNPs (Supplementary Figure 3). This is consistent with our marginal analysis in Section 2 where none of the annotations exhibited significant marginal associations with the autism z-scores. The estimated fold enrichments of FOXL1 and Nkx2-5 in the GPA analysis were 1.3 (s.e. 0.113) and 0.912 (s.e. 0.146), respectively. Both of these levels were too small to be detected with this analysis that considered only two global classes of SNPs (null and non-null).

### 4.3 PGC Schizophrenia GWAS

We applied ARoG to the SCZ2 dataset described in Section 2 with numbers of clusters K = 1, ………, 10. Best BIC values were achieved at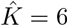 and 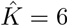 with only a 0.01% difference between the two. We carried out the rest of the analysis with 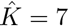

Neither ARoG(I) nor ARoG(II) selected any annotations for clusters 1-4. ARoG(I) selected FOXL1, Klf4, Prrx2, and NKX3-1 annotations for cluster

5, and Arnt::Ahr, E2F1, FOXL1, Klf4, Foxq1, Prrx2, ARID3A, and E2F4 annotations for cluster 6. Among the selected annotations, the pair of E2F1 and E2F4 and the pair of Prrx2 and ARID3A share similar sequence logos, respectively, thus, both pairs have relatively high correlations of 0.8096 and 0.6234 in the annotation score matrix. ARoG(II) retained FOXL1 and Klf4 for cluster 5 and Arnt::Ahr and FOXL1 for cluster 6 (Table 2). Overall, both cluster 5 and 6 are populated with the most genome-wide significant SNPs depicted in the Manhattan plot of Figure 1(c). Permutation testing results for ARoG(I) and ARoG(II) support significance of the selected annotations with BH adjustment at level 0.1 (Supplementary Figure 2(b)). OLS analysis of this dataset selected 15 annotations with unadjusted permutation p-values smaller than 0.05; however, none of these survived the multiple testing correction with the BH adjustment at level of 0.1. In contrast, Lasso with 5-fold cross validation tuning selected 17 annotations, of which only two (Foxq and Zfx annotations) survived the same multiple testing adjustment. The scatter plots of the SCZ2 z-scores against the ARoG(II) selected annotations exhibit associations in clusters 5 and 6 with a similar global trend across the whole SNP set (Supplementary Figure 4).

**Fig. 3:**
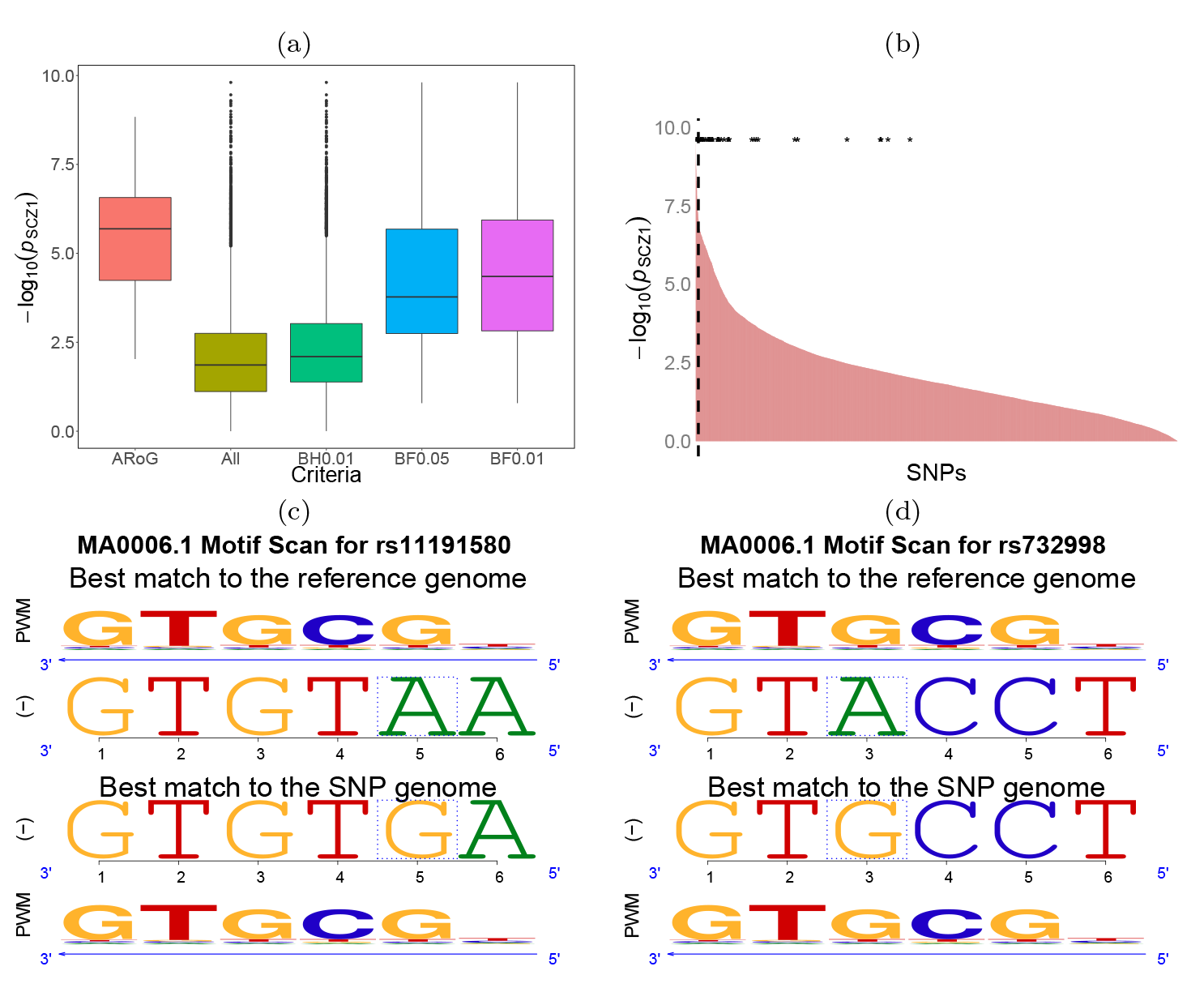
(a) SCZ1 p-values for multiple SNP sets generated based on the SCZ2 data: The SNP sets include ARoG: *ARoG SNPs*; All: All 11, 386 SNPs; BH0.01: SNPs with SCZ2 BH-adjusted p-values less than or equal to 0.01; BF0.05/BF0.01: SNPs with SCZ2 Bonferroni adjusted p-values less than or equal to 0.05/0.01. (b) SNPs ranked based on their SCZ1 significance levels. *ARoG SNPs* are marked with asterisks. The vertical dashed line depicts Bonferroni cut-off of SCZ1 analysis under significance level of 0.05. (c) Composite sequence logo of rs11191580 with the Arnt::Ahr PWM: the SNP enhances the binding of Arnt::Ahr. (d) Composite sequence logo of SNP rs732998 with the Arnt::Ahr PWM: the SNP also enhances the binding of Arnt::Ahr.

We next created a set of *ARoG SNPs* with Bonferroni corrected p-values less than 0.05 and atSNP p-values less than 0.01. While the autism dataset suffers from low power, SCZ2 dataset has many SNPs reaching genome-wide significance. Thus, we used the more stringent rule based on Bonferroni cutoff of 0.05 on the SCZ2 GWAS association p-values. *ARoG SNPs* included 14 SNPs from cluster 5 and 30 SNPs from cluster 6. Supplementary Table 1 presents genomic locations, GWAS p-values, and RegulomeDB scores [7] of these SNPs. RegulomeDB scores range from 1 for SNPs likely to affect TF binding and be linked to expression of a gene target to 7 for SNPs with no supporting data. For details on RegulomeDB scoring scheme, we refer Table 2 of Boyle et al. [7]. Of the 44 *ARoG SNPs*, fifteen have RegulomeDB score of 1, providing evidence for potential importance of these SNPs to schizophrenia. We next compared the SCZ1 association measures (p-values) of *ARoG SNPs* to those of other SNP sets one could have identified from the initial set of 11,386 SNPs without using additional functional annotation (Figure 3(a)). The other SNP sets one could define are BH0.01 (SNPs defined by BH correction at level 0. 01 on the SCZ2 p-values), BF0.05 (SNPs defined by Bonferroni correction at level 0.05 on the SCZ2 p-values), and BF0.01 (SNPs defined by Bonferroni correction at level 0.01 on the SCZ2 p-values). The intial SNP set was added as a baseline. *ARoG SNPs* are on average more significant and reproducible in the SCZ1 than the other SNP sets. Comparison of *ARoG SNPs* with a randomly selected SNP set of the same size from BF0.05 also indicated that *ARoG SNPs* are on average more significant, illustrating that the use of the functional annotation information is biasing the selection towards SNPs with reproducible associations. Figure 3(b) displays ranking of SCZ1 p-values of all 11,386 SNPs and illustrates that most *ARoG SNPs* are among the most significant SNPs with respect to SCZ1. Four of these SNPs reach genome-wide significance with Bonferroni adjustment at level 0.05 in the SCZ1 study.

**Table 2:**
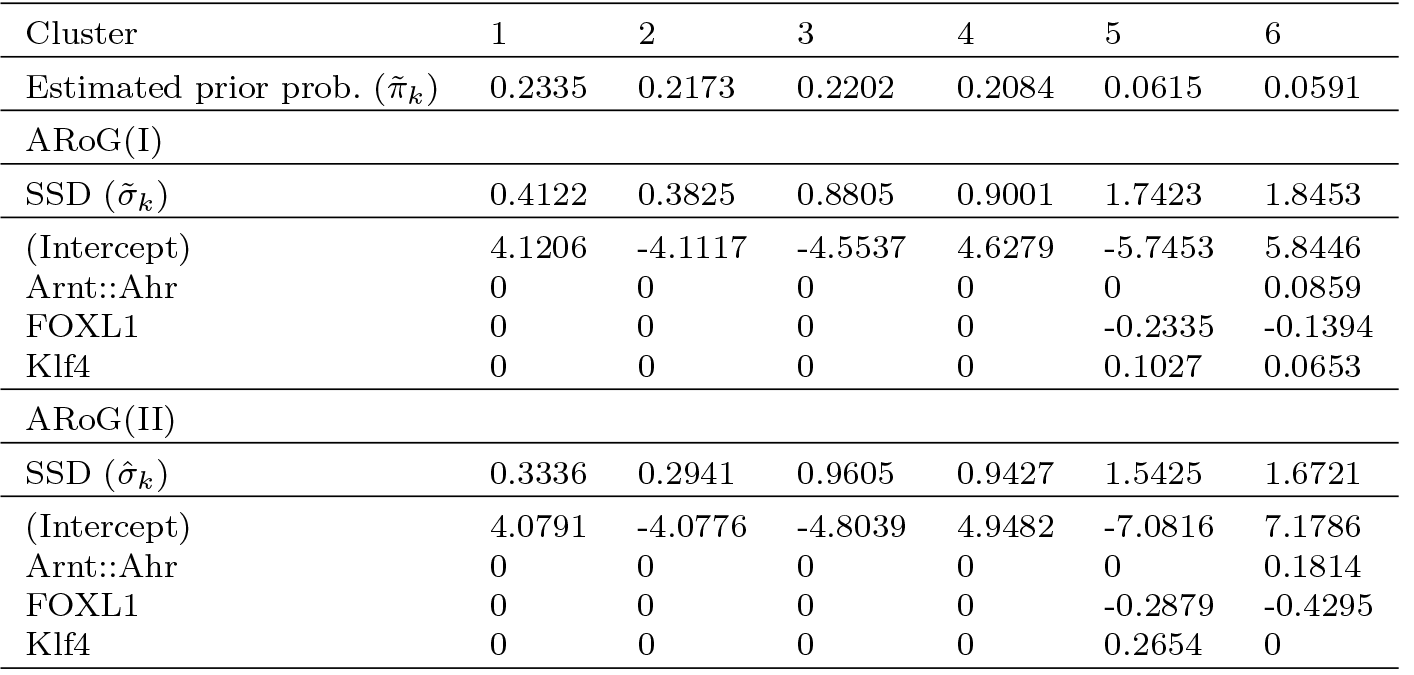
ARoG parameter estimates with PGC SCZ2 data

Next, we assessed whether any of the *ARoG SNPs* were among the schizophrenia associated SNPs from dbGaP [3]. dbGaP harbors 249 SNPs associated with schizophrenia and 42 of these are among the 11,386 SNPs we utilized. Two of the *ARoG SNPs* (rs11191580 and rs10224497, located at chr10:104,906,211 and chr7:2,149,967) are among the dbGaP SNPs. SNP rs11191580 leads to enhancement of Arnt::Ahr binding while rs10224497 seems to disrupt Arnt::Ahrbinding. Since we observed that enhanced Arnt::Ahr binding overall associated with increased schizophrenia risk in clusters 5 and 6 of the ARoG results, we further investigated rs11191580. rs11191580 is located within the 3rd intron of Nt5C2 and has rs732998, located within the 4th intron of Nt5C2, as a perfect LD partner. Their composite logo plots support that these SNPs might indeed enhance the binding of Arnt::Ahr (Figures 3(c), (d)). Furthermore, the association of rs11191580 is also validated in SCZ1 with p-value of 2.23 × 10^−8^. Although rs732998 does not quite make the genome-wide significance cut-off, it also exhibits a significant association in SCZ1 with p-value of 9.50 × 10^−8^. In summary, these two SNPs in perfect LD lead to sequence changes that are likely to improve the binding of the Arnt::Ahr complex. This complex regulates genes in response to the carcinogenic environmental contaminant 2,3,7,8– tetrachlorodibenzo-p-dioxin (TCDD). Akahoshi et al. [5] showed that Ahr TF is frequently detected in brain and over-expression of AhR causes neural differentiation of Neuro2a cells. Furthermore, recent studies support that dioxins and related chemicals influence neural development, and the AhR-signaling pathway might mediate the impact of dioxins on the nervous system [34].

Finally, as we have done for the autism dataset in Section 4.2, we applied GPA to SCZ2 dataset using all 1,175,307 SNPs that were in the dbSNP database out of the 1,179,262 genotyped SNPs. This analysis identified non-null SNPs as significantly depleted for Zfp423 annotation (Supplementary Figure 5) under Bonferroni adjusted significance level of 0.1. However, the probability that a SNP is non-null is estimated as 0.3157, thus the estimated non-null set is likely to falsely include a large number of SNPs unassociated with schizophrenia. This implies that the Zfp423 depletion is likely to be a false positive finding.

## 5 Simulation Studies

We evaluated ARoG(I) and ARoG(II) with synthetic datasets and PGC data-driven simulated datasets. As alternative methods operating on multiple annotations, we included Lasso regression, and OLS with BH correction on the regression coefficient p-values at level of 0.05. Both ARoGs enable clustering of SNPs and heterogeneous annotation coefficients across the clusters while both Lasso regression and OLS assume the homogeneity of the annotation effects. Our simluation-based evaluations focus on the detection of relevant annotations from a large set of annotations with weak effect sizes. Our application of GPA under the settings enabled by binarizing the annotation scores behaved unstably and failed in fitting the GPA model. This was mostly due to the severe sparsity of the binary annotations making their estimators be near or on the boundary of the parameter space. Therefore, we did not include GPA as an alternative method in these simulation studies.

We generated 100 simulated datasets under each scenario, where each simulated dataset consisted of training data, validation data, and test data. The validation dataset was used for selection of the optimal tuning parameter,and its sample size was increased 100 times compared to that of the training dataset. The test error was calculated as the negative log-likelihood on the test dataset with the same sample size as the training dataset. We report for each method the test error, numbers of true (TPs) and false positives (FPs), adjusted rand index (ARI) [22], receiver operating characteristic (ROC) curve, and precision recall curve. TPs and FPs for ARoG are defined by pooling selected annotations across the identified clusters. We use ARI to measure the similarity between the true SNP clusters and estimated SNP clusters. ROC curves and precision-recall curves present the performance of annotation selection in a threshold (cut-off for BH adjusted p-values for OLS, tuning parameters for Lasso and both ARoGs) agnostic manner. In these curves, we plot averages of the true positive rate (TPR), false positive rate (FPR), and precision across the 100 simulation replications.

**Table 3:**
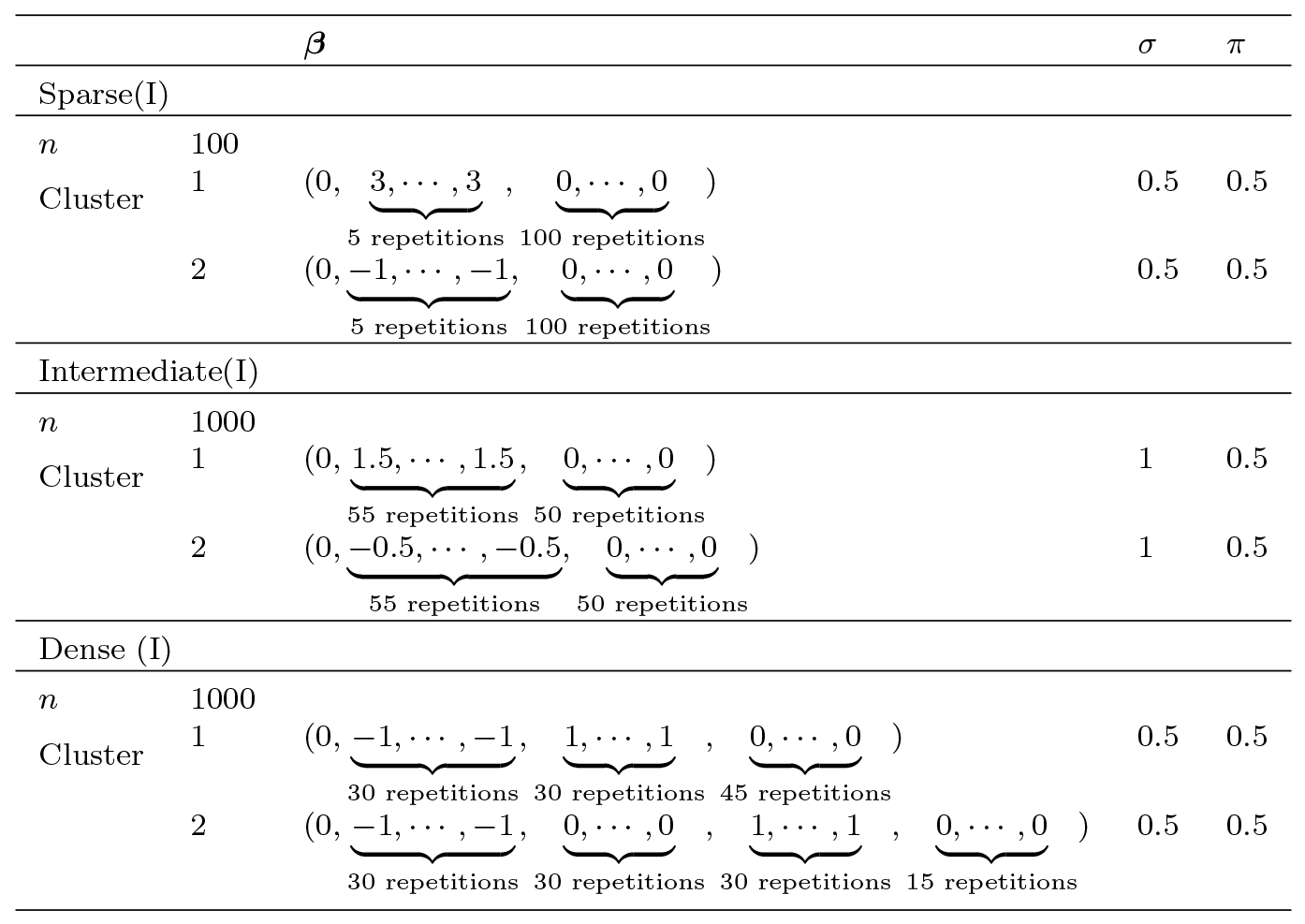
Simulation settings for Section 5.1.

| | | $\beta$ | $\sigma$ | $\pi$ |
| --- | --- | --- | --- | --- |
| Sparse(I) |  |  |  |  |
| $n$ | 100 | | | |
| Cluster | 1 | $(0, \underbrace{3, \dots, 3}_{5 \text{ repetitions}}, \underbrace{0, \dots, 0}_{100 \text{ repetitions}})$ | 0.5 | 0.5 |
| | 2 | $(0, \underbrace{-1, \dots, -1}_{5 \text{ repetitions}}, \underbrace{0, \dots, 0}_{100 \text{ repetitions}})$ | 0.5 | 0.5 |
| Intermediate(I) |  |  |  |  |
| $n$ | 1000 | | | |
| Cluster | 1 | $(0, \underbrace{1.5, \dots, 1.5}_{55 \text{ repetitions}}, \underbrace{0, \dots, 0}_{50 \text{ repetitions}})$ | 1 | 0.5 |
| | 2 | $(0, \underbrace{-0.5, \dots, -0.5}_{55 \text{ repetitions}}, \underbrace{0, \dots, 0}_{50 \text{ repetitions}})$ | 1 | 0.5 |
| Dense (I) |  |  |  |  |
| $n$ | 1000 | | | |
| Cluster | 1 | $(0, \underbrace{-1, \dots, -1}_{30 \text{ repetitions}}, \underbrace{1, \dots, 1}_{30 \text{ repetitions}}, \underbrace{0, \dots, 0}_{45 \text{ repetitions}})$ | 0.5 | 0.5 |
| | 2 | $(0, \underbrace{-1, \dots, -1}_{30 \text{ repetitions}}, \underbrace{0, \dots, 0}_{30 \text{ repetitions}}, \underbrace{1, \dots, 1}_{30 \text{ repetitions}}, \underbrace{0, \dots, 0}_{15 \text{ repetitions}})$ | 0.5 | 0.5 |

### 5.1 Synthetic Data

We generated data from several Gaussian finite mixture regression models varying the sparsity level of annotation signals as sparse, intermediate, and dense models (Table 3). The columns of the predictor matrix X are generated from an independent standard normal distribution. Supplementary Figures 6and 7 and Figure 4 present the results of these simulations.

The Sparse(I) setting is a small n, large p setting; hence, our comparisons only include ARoG(I), ARoG(II), and Lasso. Supplementary Figure 6(a) and (b) show that ARoG(I) has the smallest test error and both ARoGs have a median ARI of about 0.7. ARoG(I) and ARoG(II) have the same ARI by design since they share the same clustering assignment and Lasso has ARI of 0 since it does not perform clustering. Both versions of ARoG have high TPR with the optimal tuning parameter (Supplementary Figure 6(c)); however; ARoG(I) has an inflated FPR with an average of 10 more FPs compared to ARoG(II) (Supplementary Figure 6(d)). Lasso underselects annotations and on average has four false negatives. ROC and precision-recall curves (Supplementary Figures 6(e) and (f)) indicate that ARoGs outperform Lasso significantly. We remark that the top left corner of the ROC curves for ARoGs roughly corresponds to the results with the optimal tuning parameters presented in the boxplots of TP and FP. We also note that Lasso, ARoG(I), and ARoG(II) do not select annotations in a sequentially augmenting way as tuning parameters decrease; thus, the ROC cuves are not monotonically increasing. We also investigated this sparse setting by increasing the sample size to 1000 and observed almost perfect performance by all methods with an area under the ROC curve of 1.

Supplementary Figure 7 presents the simulation results from the intermediate models of Table 3, where almost half of the regression parameters are set as zero. The test error evaluation, ROC curves, and precision recall curves clearly indicate that ARoGs outperform OLS and Lasso. Both ARoG(I) and ARoG(II) have the optimal number of TPs, 55, with the optimal tuning parameters (Supplementary Figure 7(c)); however their numbers of FPs are substantially different (Supplementary Figure 7(d)). ARoG(II), on average, selects 2 FPs whereas ARoG(I) selects more than 45 FPs. This emphasizes the significance of the cluster-level Lasso step of ARoG(II) for reducing the numbers of FPs. ROC curves (Supplementary Figure 7(e)) reveal that although ARoG(I) and ARoG(II) overall have comparable performances, ARoG(II) performs marginally better in the top left corner, where both accurately identify all TPs, but only ARoG(II) succeeds in filtering out many FPs.

Figure 4 presents the results for the Dense (I) setting. These results agree with the superior performances of both versions of ARoG in the previous settings in terms of prediction error (Figure 4(a)). This setting also elucidates the contrast between the two ARoGs: ARoG(I) tends to select all annotations, essentially failing to achieve any variable selection whereas ARoG(II) is able to filter out FPs (Figures 4(c) and (d)). OLS tends to underselect annotations and Lasso tends to select more compared to OLS; however, still partially recovers the TPs. ARoG(II) has the best ROC and precision-recall curve performances (Figures 4(e) and (f)). This setting includes 60 annotations that are not shared between clusters in contrast to the previous settings where the annotations were shared by multiple clusters. The selection performances on these cluster-specific variables are less stable; as a result, the TPRs of both ARoGs heavily fluctuate between 0.4 and 1 in the top left corner of the ROC curves where both have small FPR. Overall, we conclude that the differences between ARoG(I) and ARoG(II) become more pronounced as the sparsity level decreases, and in such dense settings, ARoG(I) tends to have much higher FPR than ARoG(II).

**Fig. 4:**
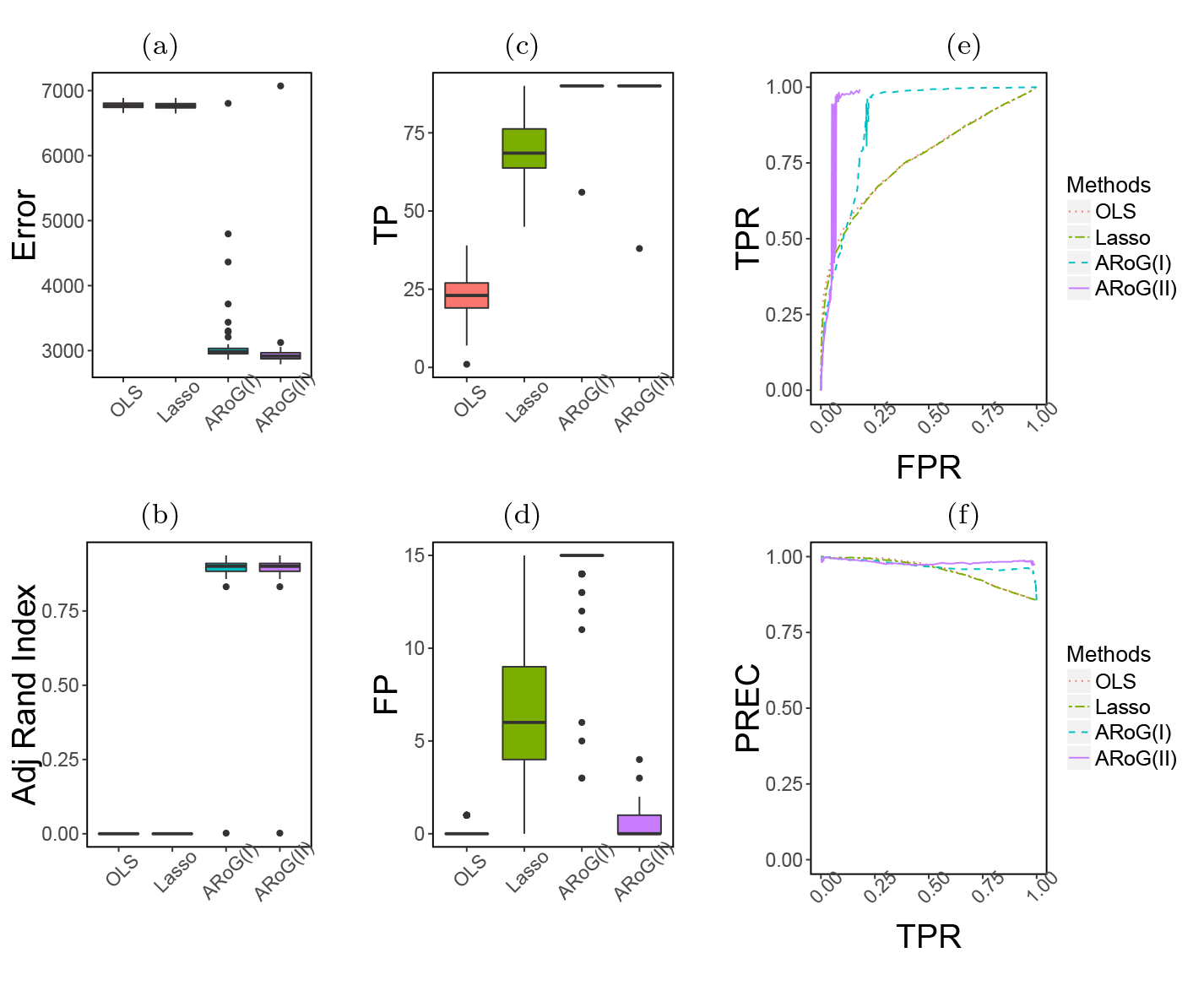
Simulation results for Dense(I) setting of Table 3.

These computational experiments involved completely simulated datasets where the predictor matrix had independent columns and was not designed to have many weak signals. Supplementary Figure 8 presents results from a weak signal setting where the actual annotation predictor matrix from the PGC autism GWAS is used to simulate data with the parameters of Supplementary Table 2. This predictor matrix has many more scores close to zero compared to the randomly generated predictor matrix in the above simulations. The overall conclusions from this setting agree well with the Sparse(I) setting.

### 5.2 PGC Analysis-Driven Data

We next evaluated the performance of ARoG in two simulation settings based on the autism and SCZ2 data analyses of Sections 4.1 and 4.3. The data were simulated based on the actual annotation score matrix from each application and the ARoG(II) parameter estimates of regression slopes, standard deviations, and prior probabilities. These data-driven simulation studies aim to capture the typical signal to noise levels observed in these type of studies.Figure 5 displays the results of the autism simulation setting. Both ARoGs reduce the prediction error compared to OLS by about 8% (Figure 5(a)). Both OLS and Lasso tend to miss TPs and thus fail to recover the underlying associations (Figure 5(c)). ARoG(I) and ARoG(II) tend to select at least one correct annotation with 86 and 78 times out of 100 repetitions, respectively. Specifically, FOXL1 is selected 71 and 61 times and Nkx2-5 is selected 49 and 42 times by ARoG(I) and ARoG(II), respectively. ARoG(II) on average filters out 2 more FPs than ARoG(I) (Figure 5(d)). Based on the ROC and precision-recall curves, ARoG(I) has the best tuning parameter-free performance followed by ARoG(II) (Figure 5(e) and (f)). OLS performs almost the same as random guess with an ROC curve on the 45 degree line.

**Fig. 5:**
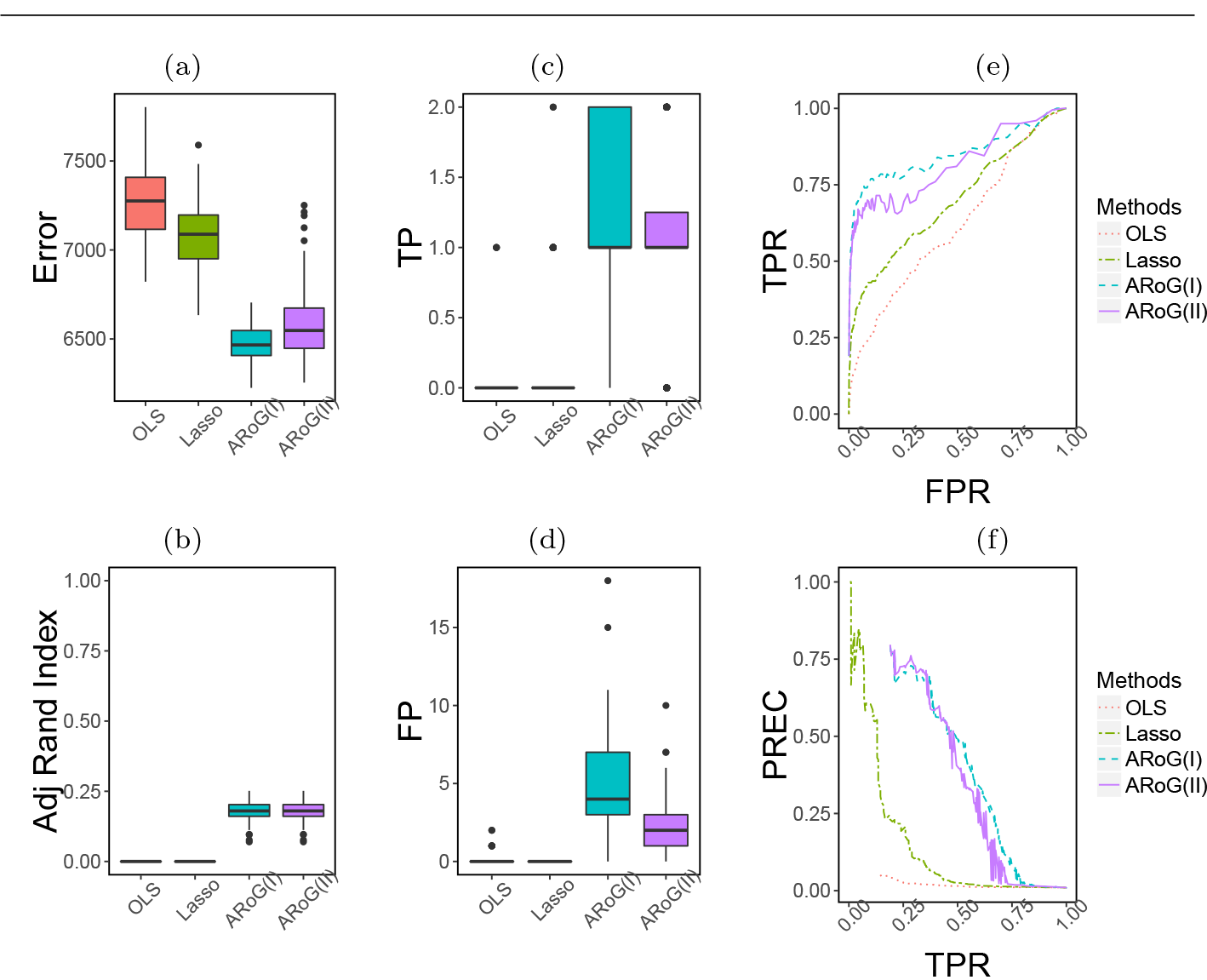
Simulation results for autism data analysis driven setting.

Supplementary Figure 9 presents the results for the SCZ2 simulation setting. Both ARoGs perform very well in terms of prediction error (Supplementary Figure 9(a)). Similar to the autism simulations, both OLS and Lasso fail to select any annotations (Supplementary Figure 9(c) and (d)). ARoG(I) and ARoG(II) tend to recover the TPs to some extent by selecting at least one correct annotation except in one simulated dataset with ARoG(II). In this setting, a trade-off between two ARoGs is clear since ARoG(I) seems better at identifying the TPs, namely, Arnt::Ahr, FOXL1, and Klf4, whereas ARoG(II) is able to more aggressively eliminate FPs. ARoG(I) has a median of 11 FPs, with more than 25 FPs in 13 of the simulated datasets. In contrast, ARoG(II) has a median of 2 FPs, with less than 10 FP annotations in almost all simulated datasets. In terms of the ROC and precision-recall curve comparisons (Supplementary Figure 9(e) and (f)), ARoG(I) exhibits a better tuning parameter-free performance compared to ARoG(II). Both OLS and Lasso perform similar to random guesses.

## 6 Discussion

We presented an integrative framework, named ARoG, for incorporating functional annotation data into GWAS analysis. The key idea behind ARoG is that even when a set of SNPs disrupts a global mechanism, e.g., pathway, that leads to disease, they might be achieving this by disrupting various submechanisms. Some might be disrupting coding sequences, some binding sites of TFs, some methylation profiles or chromatin accessibility. ARoG capitalizes on this idea and aims to identify clusters of SNPs for which GWAS association measures can be explained by a subset of functional annotations. ARoG utilizes FMRLasso [27] which enables selection among large numbers of functional annotations. We illustrated ARoG with an application to PGC data on autism and schizophrenia disorders by utilizing the impact of SNPs on TF binding affinities as functional annotations. Our analyses led to identification of SNPs which do not necessarily make the genome-wide significance cut-offs; however, are potentially worthy of following up since their GWAS associations are supplemented by their significant effects on TF binding affinities. This versatile framework provides many directions for useful extensions. First, its annotation selection capability makes it applicable with larger sets of functional annotations including TF ChIP-seq, DNase I accessibility, Histone ChIP-seq, and DNA methylation. Second, we focused our analysis on one disorder at a time; however, ARoG framework can be easily extended to simultaneously consider multiple related GWAS.

## Acknowledgements

This research was supported by National Institutes of Health grants HG007019, HG003747, and U54AI117924. The authors thank the editor and two referees for their helpful comments.

